# Long-read cDNA sequencing identifies functional pseudogenes in the human transcriptome

**DOI:** 10.1101/2021.03.29.437610

**Authors:** Robin-Lee Troskie, Yohaann Jafrani, Tim R. Mercer, Adam D. Ewing, Geoffrey J. Faulkner, Seth W. Cheetham

## Abstract

Pseudogenes are gene copies presumed to mainly be functionless relics of evolution due to acquired deleterious mutations or transcriptional silencing. When transcribed, pseudogenes may encode proteins or enact RNA-intrinsic regulatory mechanisms. However, the extent, characteristics and functional relevance of the human pseudogene transcriptome are unclear. Short-read sequencing platforms have limited power to resolve and accurately quantify pseudogene transcripts owing to the high sequence similarity of pseudogenes and their parent genes. Using deep full-length PacBio cDNA sequencing of normal human tissues and cancer cell lines, we identify here hundreds of novel transcribed pseudogenes. Pseudogene transcripts are expressed in tissue-specific patterns, exhibit complex splicing patterns and contribute to the coding sequences of known genes. We survey pseudogene transcripts encoding intact open reading frames (ORFs), representing potential unannotated protein-coding genes, and demonstrate their efficient translation in cultured cells. To assess the impact of noncoding pseudogenes on the cellular transcriptome, we delete the nucleus-enriched pseudogene PDCL3P4 transcript from HAP1 cells and observe hundreds of perturbed genes. This study highlights pseudogenes as a complex and dynamic component of the transcriptional landscape underpinning human biology and disease.

## Background

Pseudogenes are gene copies which are thought to be defective due to frame-disrupting mutations or transcriptional silencing [1,2]. Most human pseudogenes (72%) are derived from retrotransposition of processed mRNAs, mediated by proteins encoded by the LINE-1 retrotransposon [3,4]. Due to the loss of parental *cis*-regulatory elements, processed pseudogenes were initally presumed to be transcriptionally silent [1] and were excluded from genome-wide functional screens and most transcriptome analyses [2]. Transcriptomic surveys of cancer [5] and normal human tissues [6] by high-throughput short-read sequencing suggest that pseudogene transcription may be widespread. However, studies of pseudogene transcription are hindered by the limited capacity of short-read sequencing, and microarray hybridisation, to discriminate pseudogenes from their highly similar parent genes [2,7]. Most full-length pseudogene transcripts found to date were identified by relatively low-throughput capillary sequencing of full-length cDNA libraries [8–10]. As a result, the extent of the human pseudogene transcriptome in most spatiotemporal contexts remains largely unresolved.

Pseudogene transcripts can control the expression of their parent genes by acting as competitive endogenous RNAs [11] (ceRNAs), antisense transcripts [12], precursors for small interfering RNAs [13,14] (siRNAs) and piwi-interacting RNAs [15] (piRNAs). Whilst most pseudogenes are presumed to act by noncoding mechanisms, some retain the capacity to encode full-length or truncated proteins [16–19].

## Results and discussion

Long-read cDNA sequencing via Pacific Biosciences Isoform Sequencing (PacBio Iso-Seq) or Oxford Nanopore Technologies is a potentially powerful approach to identify full-length pseudogene transcripts and accurately differentiate pseudogenes and their parent mRNAs. PacBio Iso-Seq is particularly suitable for this application due to the high consensus accuracy enabled by circular consensus reads. To comprehensively survey the human processed pseudogene transcriptome, we sequenced high quality RNA from 20 normal mixed adult and foetal human tissues (Qiagen XpressRef Universal Total RNA) on a Sequel II platform **(Fig. 1a)**. To further broaden the biological scope of our analysis, we integrated our data with a deep PacBio in-house Sequel II dataset of 6,775,127 full-length reads from a mixture of 10 human cell lines [20]. We aligned the reads to the human reference genome (hg38) at high stringency (q60) and compared the identified transcript isoforms to Gencode [21] annotations using SQANTI2, a bioinformatics QC tool designed to annotate full-length transcript (Iso-Seq) data with respect to a reference transcriptome [22].

**Fig. 1.**
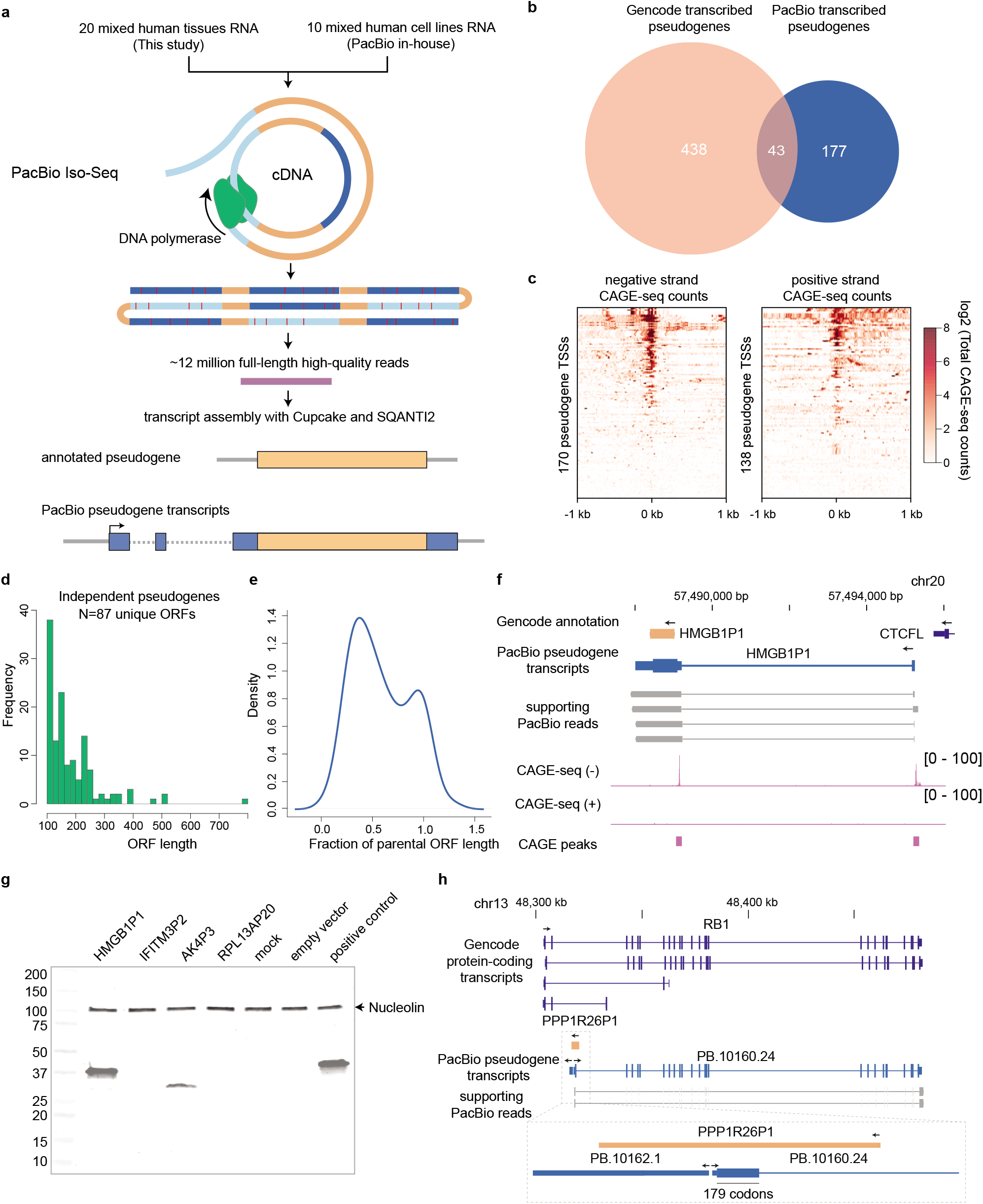
Long-read cDNA sequencing elucidates the human pseudogene transcriptome. **a** Full-length consensus PacBio cDNA reads from normal tissues and cell lines were compared to Gencode annotations to generate a pseudogene transcriptome. **b** Most transcribed pseudogenes identified here were absent from Gencode. **c** The transcription start sites (TSSs) of full-length pseudogene transcripts are enriched for CAGE-seq signal (data from FANTOM5 [24]). **d** ORF lengths of potentially coding-independent pseudogene transcripts. **e** Fraction of parental ORF length found intact in transcribed potentially coding-independent pseudogenes. **f** HMGB1P1 has a novel 5’ exon and is transcribed from an upstream CAGE-confirmed TSS. **g** Expression of 3XHA-tagged pseudogene ORFs in HEK293T cells detected by western blot. HMGB1P1 and AK4P3 are translated when expressed in cultured cells. **h** A novel isoform of retinoblastoma (RB1) is transcribed from a TSS located within the pseudogene PPP1R26P1. The pseudogene sequence adds 179 codons to the RB1 ORF.

We identified 1170 polyadenylated transcripts, each supported by at least two full-length reads, that overlapped 521 processed pseudogenes (**Table S1**). 220 pseudogenes (318 transcripts) transcribed in sense (the same orientation as their parent gene) were independent (non-intronic and have greater overlap with the pseudogene than other gene models) of known genes, only 43 of which were previously annotated as transcribed pseudogenes in Gencode (**Fig. 1b,** for examples see **Fig. S1**). 101 of these transcripts were multi-exonic, and the vast majority (84%) did not incorporate splice junctions with known Gencode transcripts. Pseudogenes are typically transcribed in the same orientation as their parent genes [23]. However, we found 78/396 (20%) of independent pseudogene transcripts were produced in antisense with respect to their parent gene (for examples see **Fig. S2a-b**). In contrast only 68/2669 (6.3%) of unprocessed pseudogenes are transcribed in antisense (**Fig S2c**). This difference may be attributable either to the propensity for novel downstream promoter elements to regulate expression of a retrotransposed pseudogene relative to unprocessed pseudogenes (which retain their parental promoter) or to selection for regulatory potential. Manual inspection of the antisense pseudogene TSSs did not reveal an obvious initiation site bias. Antisense pseudogene transcripts have significant potential to regulate their parent genes by antisense-mediated translational inhibition or by processing into siRNAs [12–14]. In support of the pseudogene transcripts identified here being full-length, we intersected our data with an atlas of Cap Analysis Gene Expression 5’ mRNA sequencing (CAGE-seq) data generated by FANTOM5 [24]. CAGE signal was highly enriched at pseudogene transcription start sites (TSSs) (**Fig. 1c**) and 41% of pseudogene TSSs were within 100bp of a FANTOM5 [24] CAGE peak, indicating that a large fraction of pseudogene transcripts have accurate 5’ ends. The proportion of pseudogene transcripts that overlap CAGE peaks is lower than for protein-coding transcripts (69%) and comparable to lincRNA transcripts (44%) (**Fig. S3**). 51% of independent pseudogene transcripts were supported by the cutoff of two full-length reads (**Fig. S4**), and our datasets do not comprehensively capture diversity of human cell-types and developmental stages. Therefore it is probable that our data still significantly underestimate pseudogene transcription; further transcripts would very likely be identified by increased sequencing depth applied to individual tissues or cell types.

Next, we annotated the coding potential of independent pseudogene transcripts using SQANTI2. 160/318 pseudogene transcripts (50%) encode putative proteins that are >100 amino acids in length (**Fig. 1d)** and, strikingly, 53 of the pseudogene open reading frames (ORFs) were >90% of the length of the parent gene ORF (**Fig. 1e**). An illustrative example of a potentially coding pseudogene transcript is the processed pseudogene of the high mobility group box 1 on chromosome 20 (HMGB1P1). The Gencode HMGB1P1 annotation is a single contiguous region of 98% identity to the HMGB1 ORF, which has no introns (**Fig. 1f)**. Iso-Seq revealed that HMGB1P1 was transcribed from an upstream promoter, which yields a novel 5’ exon, and was supported by a FANTOM5 CAGE peak. HMGB1P1 contained no frameshift mutations and encoded a protein of the same length as HMGB1, with an intact HMG domain. To assess the coding potential of HMGB1P1 and other pseudogene transcripts, we amplified the 5’ exons and coding sequence of four spliced pseudogenes with intact ORFs (HMGB1P1, AK4P3, IFITM3P2 and RPL13AP20) and cloned them into a vector with a C-terminus 3XHA tag. Transfection into HEK293T cells resulted in clear translation of the HMGB1P1 and AK4P3 transcripts (**Fig. 1g, Fig. S5)**. To further substantiate that pseudogenes can be translated *in vivo* we interrogated the neXtprot human proteomics database [25]. Eleven potentially-coding pseudogenes have entries in neXtprot of which four, HMGB1P1, SUMO1P1, MSL3P1 and PLEKHA8P1 have matched unique peptides (**Fig. S6**). Thus, pseudogene transcripts can encode intact proteins that are translated in human cells.

To determine if the pseudogene ORFs are subject to purifying selection we identified orthologous positions in non-human primate genomes by aligning the human transcripts with a 1000bp window on each side to higher primates (chimp, gorilla, orangutan, and rhesus). The alignments were then further refined to identify orthologous cDNA sequences and the resulting ORFs were translated into amino acid sequences (Methods). The extent of selection on pseudogene ORFs was determined by maximum likelihood estimation of the ratio of substitution rates between two divergent species that result in nonsynonymous vs synonymous changes (dN/dS). A ratio of >1 suggests diversifying selection whilst a ratio of <1 is consistent with purifying selection. This index has been used as evidence of conserved function for human processed pseudogenes [26]. Of the 32 pseudogene ORFs which are conserved in Rhesus Macaque and have sufficient nucleotide diversity 29/35 (85%) have a dN/dS <1 (median 0.4483) suggesting that most of the conserved pseudogene ORFs were under purifying selection across 25M years of evolution (**Table S2**).

In addition to independent protein-coding potential, pseudogenes can contribute to the coding sequences of known genes. We found that 93 protein-coding genes contained coding sequences derived from pseudogenes, often adding hundreds of codons (**Fig. S7**). Notably, the pseudogene PPP1R26P1 constitutes most of a novel 5’ exon fused to the major tumour suppressor gene retinoblastoma (RB1), adding 179 codons to RB1 from the antisense strand of PPP1R26P1 (**Fig. 1h)**. Indeed, PPP1R26P1 was previously shown to constitute an alternative imprinted RB1 promoter [27]. Gene-pseudogene fusion transcripts can add novel domains to genes, such as the case of HMGN1P18, which adds a HMGN domain to CPED1 (**Fig. S8a)**. Even well-characterised long noncoding RNAs (lncRNAs) may have isoforms that encode pseudogene proteins, including the lncRNA FIRRE [28], which has an isoform that splices into an intact MCRIP2 pseudogene (**Fig. S8b**).

To better evaluate spatial patterns of pseudogene transcription, we aligned deep RNA-sequencing from 16 adult tissues of the Illumina Body Map 2.0 [29] to our independent pseudogene annotations (**Table S3**). PacBio-identified pseudogene transcripts were highly tissue-specific (**Fig. S9)**, and divergent from expression of their parent genes. For example, AMD1P4 and YWHAEP1 expression is, respectively, liver- and testis-specific (**Fig. 10a-b**) whilst their parent genes are broadly expressed (**Fig. S10c-d**), indicating that pseudogene expression is controlled by distinct regulatory elements. Short-read sequencing can therefore be leveraged to quantify the expression of pseudogene transcripts discovered by long-read sequencing.

To determine if long-read sequencing outperforms short-read sequencing at assembling pseudogene transcripts we generated matched PacBio and Illumina datasets from the haploid leukaemia cell line HAP1. Without a reference transcriptome, 71% of the 163 HAP1 PacBio pseudogene transcripts were detected by short-read transcript assembly with StringTie [30], whilst with Gencode (v29) as a reference, 91% of pseudogene transcripts were detected. However, the reference-guided short-read assembled transcripts were significantly shorter than the PacBio transcripts (average of 750bp shorter, p=8.5E-11 Mann-Whitney) indicating that these assemblies do not cover complete transcripts (**Fig. S11a-b**).

Transcribed pseudogenes can regulate gene expression through coding-independent mechanisms. Haploid cells are ideally suited to genetic manipulation as only a single allele needs to be inactivated for complete loss-of-function [31,32]. Among 163 independent pseudogene transcripts, we identified PDCL3P4 as being highly expressed from a human endogenous retrovirus-K (HERV-K) long terminal repeat (LTR) promoter on chromosome 3 **(Fig. 2a)**. PDCL3P4 is derived from retrotransposition of phosducin-like 3 (PDCL3), a putative chaperone protein implicated in angiogenesis and proliferation [33]. Unlike the well-characterised pseudogene PTENP1 [11,34] (**Fig. S12a**), expression of PDCL3P4 expression does not correlate with that of its parent gene, indicating they are likely regulated independently (**Fig. S12b**). As a route to test the regulatory impact of the PDCL3P4 locus, we deleted the pseudogene from HAP1 cells with CRISPR-Cas9 genome editing by directing a Cas9 endonuclease-guide RNA (gRNA) complex to unique genomic regions flanking PDCL3P4. Three independent clonal PDCL3P4 knockout lines were generated with a combination of gRNAs to reduce the risk of off-target mutations. Genotyping of the PDCL3P4 locus in knockout cells revealed that two of the lines, null1 and null3, contained complete deletions (**Fig. 2b**) whilst the remaining line (null2) contained a complete deletion and a 154bp insertion at the site of CRISPR-Cas9 mutagenesis. Replicate RNA-seq confirmed PDCL3P4 expression was entirely abrogated in each knockout line (**Fig. 2c,** null1:N=2, null2:N=4, null3: N=4,). Additionally, 137 differentially expressed genes (DEGs) were detected in the three knockout lines, compared to wild-type cells (N=4) and a control clone (N=3) in which PDCL3P4 deletion was unsuccessful (FDR=0.01, **Table S4**) (**Fig. 2d**). Changes in gene expression were consistent between the independent clonal knockouts, indicating that these expression changes do not represent off-targets. PDCL3 expression was unaffected in the knockout lines, as were any genes within 200kb of PDCL3P4. Perturbation of unannotated *cis*-regulatory elements within PDCL3P4 was therefore unlikely to drive downstream differential gene expression, although this does not fully exclude the potential caveat of CRISPR-Cas9 genetic manipulation otherwise impacting transcription. The PDCL3P4 ORF was highly disrupted, suggesting the pseudogene could act as a lncRNA. Indeed, PDCL3P4 transcripts were enriched 5.17-fold in the nucleus compared to the cytoplasm in wild-type HAP1 cells (**Fig. 2e**), consistent with the subcellular localisation of a large fraction of lncRNAs to the nucleus [35–37]. Therefore, transcribed noncoding pseudogenes may impact the transcriptome in a parent gene-independent manner.

**Fig. 2.**
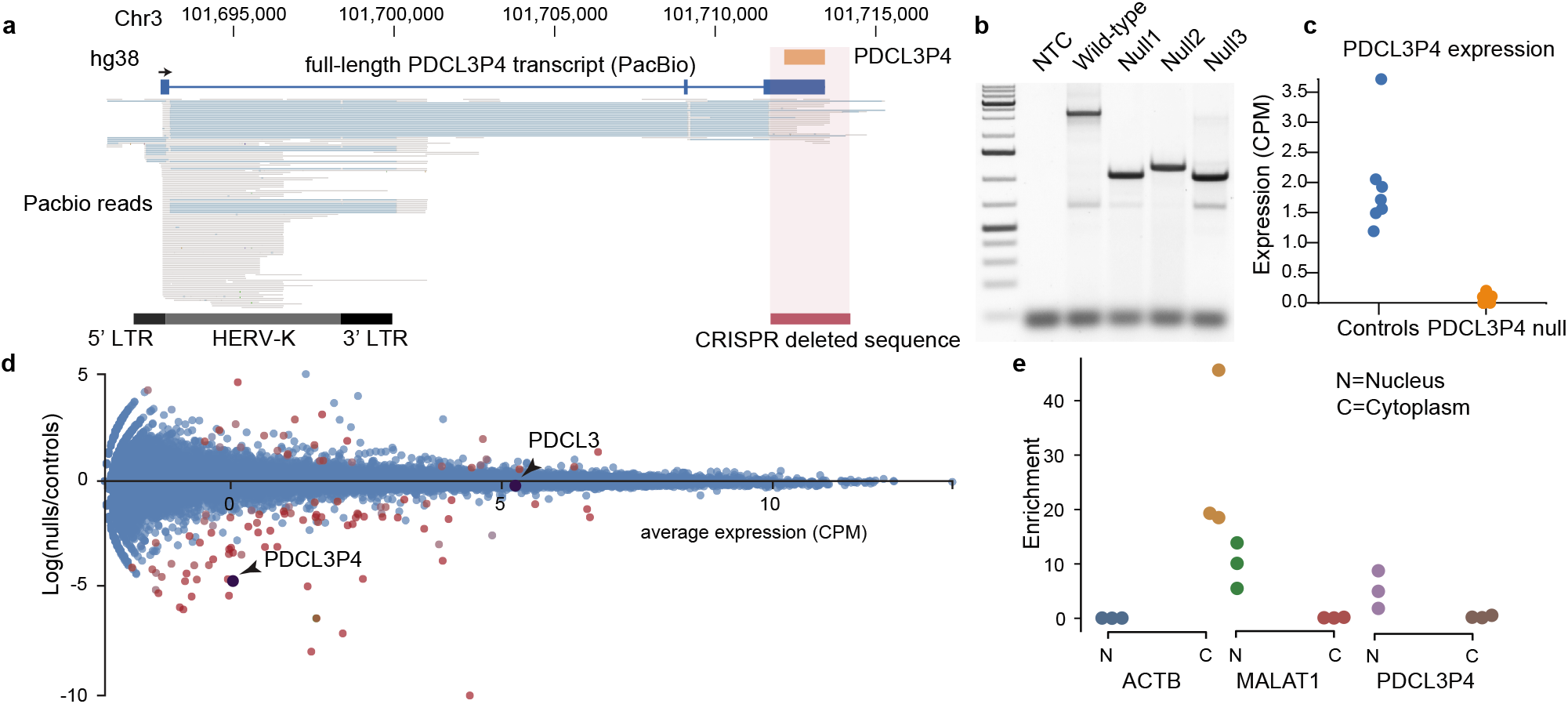
Deletion of PDCL3P4 impacts the transcriptome of haploid cells. **a** PDCL3P4 is a pseudogene transcribed in HAP1 cells from the canonical long terminal repeat (LTR) promoter of an upstream human endogenous retrovirus-K (HERV-K) sequence. Grey bars within the PacBio reads represent exons and light blue bars represent introns. **b** CRISPR-Cas9 genome engineering removes the retroposed portion of PDCL3P4 from the HAP1 genome in three independent clones. **c** PDCL3P4 expression is ablated in PDCL3P4 mutant clones. **d** PDCL3P4 ablation disrupted the expression of more than 137 genes, while PDCL3 expression was unaffected. **e** PDCL3P4 transcripts were enriched in the nucleus. The nuclear-localised noncoding RNA MALAT1 and mRNA ACTB act as controls.

## Conclusions

Here, we define a complex tissue-specific pseudogene transcriptome using PacBio long-read sequencing, validated by orthogonal CAGE-seq and RNA-seq datasets. This high-quality annotation can be utilised as a resource for transcriptomic analyses and to design functional screens. We contribute to the growing body of evidence that pseudogene translation may be widespread [38–41] and provide proof-of-principle evidence that noncoding pseudogenes can regulate the cellular transcriptome by mechanisms independent of their parent gene. Future work will elucidate the mechanism through which PDCL3P4 affects gene expression. Collectively these data suggest pseudogenes have prematurely been assumed to be functionless and numerous annotated pseudogenes produce protein-coding or noncoding transcripts. This study is a foundation for the use of long-read transcriptome sequencing to comprehensively identify full-length pseudogene transcripts and thereby better understand a major and underappreciated component of the transcriptional landscape, and its impact on human biology and pathology.

## Methods

### PacBio sequencing

XpressRef Universal Total RNA (Qiagen cat # 338112) and HAP1 (Horizon) RNA (RIN 10) was sequenced on the Pacific Biosciences Sequel II platform at the University of Maryland Institute for Genome Sciences. Libraries were prepared using the PacBio Iso-Seq library preparation, which amplifies full-length polyadenylated transcripts without size selection. Two 8M flow cells were used to generate the Universal Total RNA data (6,002,282 full-length reads) and one 8M flow cell was used for the HAP1 data (4,098,069 full-length reads).

### PacBio data processing

Full-length circular consensus reads were identified with lima v *1.10.0* (https://github.com/PacificBiosciences/barcoding) using the settings lima --isoseq --dump-clips --no-pbi --peek-guess -j 24 removing the primer sequences:

>NEB_5p GCAATGAAGTCGCAGGGTTGGG
>NEB_Clontech_3p GTACTCTGCGTTGATACCACTGCTT

Transcripts were refined and polyA tails removed with isoseq3 v3.2.2 (https://github.com/PacificBiosciences/IsoSeq) *refine* --require-polya. Full-length reads were converted to fastq format using bamtools v2.5.1 [42] *convert*. High quality clustered reads were aligned with minimap2 [43] v2.17-r941 with the settings -ax splice --secondary=no -C5 -O6,24 -B4 -uf. Redundant isoforms were collapsed with cDNA Cupcake (https://github.com/Magdoll/cDNA_Cupcake) 9.1.1 collapse_isoforms_by_sam.py --dun-merge-5-shorter. 5’ degraded transcripts were removed with filter_away_subset.py and transcript abundance counted with get_abundance_post_collapse.py. SQANTI2 v6.0.0 (https://github.com/Magdoll/SQANTI2) was used to classify the high quality clustered non-redundant reads with respect to Gencode [21] (v29) and FANTOM5 [24] CAGE peaks.

### Identification of pseudogene transcripts

Pseudogene transcripts were identified by intersecting the PacBio transcripts with Gencode pseudogenes [23] (v29) using bedtools v2.29.2 [44]. To confirm that these transcripts intersected directly with a retrotransposed copy (rather than with another exon of spliced transcript that is annotated as a transcribed pseudogene) we further intersected these transcripts with the retrogenes.v9 [10] track downloaded from the UCSC genome browser [45]. Independent pseudogene transcripts were classified as those assigned the name of a Gencode pseudogene by SQANTI2.

### Illumina Human BodyMap quantification

Sequence data in .fastq format was obtained from the Illumina Human BodyMap 2.0 Project [29] (SRA accession PRJNA144517) and aligned to the Ensembl GRCh38 primary assembly with STAR 2.7.3a [46]. The STAR reference was built using Ensembl build 101 [47] with the independent pseudogene models added to the gtf file. Reads were aligned with default parameters with the exception of --outFilterMultimapNmax 1 to limit multi-mapping reads. Reads were counted against the independent pseudogene model using htseq-count 0.11.2 [48]. Single-end and paired-end reads were counted separately across sequence runs and summed into a single count per-tissue. TMM-normalised cpm values were produced using edgeR 3.24.3 [49] and transformed to log2(cpm+1). Read mappings were visualised in IGV [50].

### Conservation analysis

Conservation was assessed by aligning each pseudogene transcript and a 1000bp flank on both ends to the human (hg38), chimp (panTro6), gorilla (gorGor6), orangutan (ponAbe3), and rhesus (rheMac10) genome assemblies using BLAT [51] (gfServer -stepSize=5). The human cDNA (derived from “CDS” entries in the input .gff file) was aligned within the larger transcribed region using exonerate [52] in “cdna2genome” mode, and the highest scoring alignment was presumed to be the ortholog of the human cDNA. Where the cDNA sequence length was divisible by 3, the ORFs were translated and compared to the human ORF using exonerate in “ungapped” mode. For each transcript that contained at least one CDS and aligned to two or more species, pairwise dN/dS statistics were obtained as follows. Multiple sequence alignments of the ORF protein sequences were performed via clustal omega [53] with default parameters, cDNAs were codon-aligned using PAL2NAL [54], and pairwise dN/dS was computed via codeml from PAML 4.9j [55]. A python script for carrying out these methods is available at https://gist.github.com/adamewing/3a4cfa8eb1a333ee9c497538ce30b6db.

### Cell culture

Low passage (**≤** p10) HAP1 cells (Horizon) were cultured in Iscove’s Modified Dulbecco’s Medium (IMDM) (Gibco cat # 12440-053) supplemented with 10% Fetal Bovine Serum (Sigma Aldrich cat # F2442) and 1% Penicillin / Streptomycin (Gibco cat # 15140-122) and grown in a tissue culture incubator (37°C, 5% CO2). Cells were not maintained above p18. HEK293T cells (ATCC) were cultured in Dulbecco’s Modified Eagle Medium (DMEM) (Gibco cat # 21969035) supplemented with 10% Fetal Bovine Serum (Sigma Aldrich cat # F2442) and 1% Penicillin / Streptomycin (Gibco cat # 15140-122) and grown in a tissue culture incubator (37°C, 5% CO2).

### Custom guide RNA design

Genomic DNA flanking the PDCL3P4 locus was examined for evidence of enhancer marks or transcriptional activity using the GeneHancer and Layered H3K27Ac tracks on the UCSC Genome Browser. Approximately 700bp of up- and downstream genomic sequence without evidence of functional activity was selected to design custom CRISPR-Cas9 gRNAs using the IDT Custom Alt-R^®^ CRISPR-Cas9 Guide RNA Design Tool (https://sg.idtdna.com/site/order/designtool/index/CRISPR_SEQUENCE). Two upstream and two downstream gRNAs were chosen based on optimal on- and off-target scores as well as by manual inspection of off-target hits to corresponding gRNA design.

### CRISPR-Cas9 genome engineering

PDCL3P4 knockout lines were generated in HAP1 cells (Horizon) following the Alt-R CRISPR-Cas9 System: Cationic lipid delivery of CRISPR ribonucleoprotein complexes into mammalian cells protocol (IDT). Pools of low passage HAP1 cells were individually reverse transfected with alternating combinations of upstream and downstream gRNA (**Table S5**): ribonucleoprotein (RNP) complexes labelled with a fluorescent dye (ATTO-550) using Lipofectamine CRISPRMAX Transfection Reagent (Thermo Fisher Scientific cat # CMAX00008).

Cells were incubated with the transfection complexes in a tissue culture incubator (37°C, 5% CO2) for 48h and then prepared for fluorescence-activated cell sorting (FACS). Cells were stained using LIVE/DEAD^®^ Fixable Aqua Dead Cell Stain (Thermo Fisher Scientific cat # L34966) following the manufacturer’s instructions and then resuspended in Hank’s Balanced Salt Solution (HBSS) (Thermo Fisher Scientific cat # 14025076). Cell populations were gated on the BD FACSAria™ Fusion Sorter based on viability and a positive signal for ATTO-550. Single cells were sorted into individual wells in a 96-well tissue culture plate containing Iscove’s Modified Dulbecco’s Medium (IMDM) (Thermo Fisher Scientific cat # 12440053). Single cells were clonally expanded and genomic DNA was extracted from half the clonal population using QuickExtract™DNA Extraction Solution (Epicentre cat # QE09050) following the manufacturer’s instructions. Null1 and null2 were derived using upstream gRNA2 and downstream gRNA1, whilst null3 was generated using upstream gRNA2 and downstream gRNA2. The control clone unsuccessful for PDCL3P4 deletion was treated with upstream gRNA2 and downstream gRNA1.

### Sanger sequencing

Individual clones were assessed for PDCL3P4 knockout by performing a genotyping PCR with Q5^®^ High-Fidelity DNA Polymerase (New England BioLabs cat # M0492S) and primers placed outside of the gRNA cut sites. PCR products were run on a 1% agarose gel to inspect for the presence of a wild-type or knockout amplicon. Amplicons indicative of PDCL3P4 knockout were cut from the agarose gel and DNA extracted using the QIAquick Gel Extraction Kit (Qiagen cat # 28704). DNA was capillary sequenced by the Australian Genome Research Facility (AGRF) to validate the knockout.

### Quantitative real-time PCR

RNA was extracted from validated clones and wild-type HAP1 cells using the RNeasy Mini Kit (Qiagen cat # 74104) following the manufacturer’s instructions and then treated with TURBO DNA-free™ Kit (Life Technologies cat # AM1907) to remove genomic DNA. DNA-free RNA was used for quantitative real-time PCR (qRT-PCR) to validate the PDCL3P4 knockout. Primers for PDCL3P4 were designed to target a SNP-containing region to mitigate off-target binding to the parent gene. Both a standard curve and melt curve were performed using *Power* SYBR^®^ Green RNA-to-CT™ 1-Step Kit (Thermo Fisher Scientific cat # 4389986) to ensure optimal amplification efficiency and specificity of the primers. qRT-PCR was performed on RNA extracted from validated clones and wild-type HAP1 cells using the abovementioned kit on the Applied Biosystems ViiA™ 7 Real-Time PCR machine, and Ct values were normalised to ACTB expression. PDCL3P4 expression was compared between wild-type HAP1 cells and the knockout clones to validate the absence of expression in knockout clones (data not shown).

### RNA-seq

The three validated PDCL3P4 knockout clones, wild-type HAP1 cells and a clone unsuccessful for the knockout were then prepared for RNA-seq (20M PE150 reads) (Novogene). Cells from each condition (knockout, wild-type, unsuccessful knockout) were seeded in five replicates in a 6-well tissue culture plate and left to proliferate for 24h. RNA was extracted from each well and genomic DNA removed using the abovementioned kits. A DNA contamination PCR was performed using MyTaq™ DNA Polymerase (Bioline cat # BIO-21105) and primers for ACTB to check for the presence of genomic DNA. RNA quality from DNA-free samples was measured on the Agilent 2100 Bioanalyser (Agilent cat # G2939BA) using the Agilent RNA 6000 Nano Kit (Agilent cat # 5067-1511). RNA concentration and purity were measured on the NanoDrop™ Lite Spectrophotometer (Thermo Fisher Scientific cat # ND-LITE-PR). The four RNA samples per condition with the highest RIN scores and purity (A260/280 ratio) were prepared for shipment. 2μg of RNA was dried in RNAstable^®^ tubes (Sigma Aldrich cat # 93221-001-1KT) following manufacturer’s instructions and heat sealed in a desiccant bag. Two out of four replicates for null1 and one out of four replicates for the control clone unsuccessful for PDCL3P4 knockout did not pass QC before library preparation. All other samples and replicates passed QC and underwent library preparation for sequencing. RNA-seq data was analysed using STAR 2.7.3a [46] to align RNA-seq reads to GRCh38, htseq-count 0.11.2 [48] to quantify read counts against Ensembl genes GRCh38.83 [47], and EdgeR [49] for DEG analysis via Degust [56]. Wild-type and control clone replicates were compared against the three null line replicates collectively. Normalised read counts for each sample are in **Table S4.** The most differentially expressed genes are consistently different between nulls compared to either wild-type cells or to the clone in which excision was unsuccessful.

### Cloning

cDNA was generated from 5μg of Human XpressRef Universal Total RNA (Qiagen cat # 338112) using Superscript III Reverse Transcriptase (Invitrogen cat # 18080093) following manufacturer’s instructions. HMGB1P1, IFITM3P2, AK4P3 and RPL13AP20 were amplified from cDNA with Q5^®^ High-Fidelity DNA Polymerase (New England Biolabs cat # M0492S) using primers that amplify the novel full-length transcript and contain HindIII and NotI restriction sites (**Table S5**). PCR amplicons were run on a 1% agarose gel and then purified using the QIAquick Gel Extraction kit (Qiagen cat # 28706). 1μg of DNA was digested with HindIII (New England Biolabs cat # R0104S) and NotI (New England Biolabs cat # R0189S) for 2h at 37°C. Digested PCR products were cleaned up using the QIAquick PCR Purification kit (Qiagen cat # 28104) and then ligated into pcDNA 3.1 3xHA cut with HindIII and NotI using the Quick Ligation kit (New England Biolabs cat # M2200L) following manufacturer’s instructions. Plasmid constructs were transformed into One Shot™ TOP10 Chemically Competent *E. coli* (Invitrogen cat # C404003) and plated on agar plates containing ampicillin 100mg/mL (Sigma-Aldrich cat # A1593). Several colonies were cultured overnight in 5mL of luria broth and plasmid DNA was extracted using the QIAprep Spin MiniPrep kit (Qiagen cat # 27106). Plasmid DNA was sent for sequencing at the Australian Genome Research Facility (AGRF) to confirm the presence of the full-length pseudogene transcripts. Confirmed plasmids were cultured overnight in 50mL of luria broth and then DNA extracted using the QIAGEN Plasmid *Plus* Midi kit (Qiagen cat # 12945).

### Western blot

HEK293T cells (ATCC^®^ CRL-3216™) were seeded at a density of 5×10^5^ cells/well in a 6-well culture plate (Sigma Aldrich, cat # CLS3516). The following day, cells were transfected in a 3:1 ratio of FuGENE^®^ HD Transfection Reagent (Promega cat # E2311) and plasmid DNA (pcDNA3.1-HMGB1P1-3xHA, pcDNA3.1-IFITM3P2-3xHA, pcDNA3.1-AK4P3-3xHA, pcDNA3.1-RPL13AP20-3xHA) in OptiMEM (Thermo Fisher cat # 31985062). Empty pcDNA3.1-3xHA was used as a negative control, whilst pcDNA3.1-3xHA-TurboID [57] served as a positive control. Cells were incubated with the transfection complex in a tissue culture incubator (37°C, 5% CO2) and after 24h protein lysate was extracted using RIPA Lysis and Extraction Buffer (Thermo Fisher cat # 89900) containing cOmplete™ Protease Inhibitor Cocktail (Sigma Aldrich cat # 4693116001). Protein concentration was measured using the Pierce™ BCA Protein Assay Kit on the POLARstar^®^ Omega microplate reader (BMG Labtech). 7μg of protein lysate was diluted with 4x Laemmli Sample Buffer (BioRad cat # 1610747) containing 10% 2-Mercaptoethanol (Sigma Aldrich cat # M6250) and reduced for 5min at 98°C. Samples were loaded into a 4–20% Mini-PROTEAN^®^ TGX™ Precast Protein Gel (BioRad cat # 4561094) and run in a Mini-PROTEAN Tetra Vertical Electrophoresis Cell for 35min at 200V. Proteins were transferred onto the iBlot™ Transfer Stack (Thermo Fisher cat # IB301001) using the iBlot™ Gel Transfer Device (Thermo Fisher) 7min program. The membrane was dried overnight and activated in 1xTBS for 5min and then blocked in Odyssey Blocking Buffer TBS (LI-COR cat # 927-50000) for 1h at room temp. The membrane was incubated with purified anti-HA.11 epitope tag antibody (BioLegend cat # 901503) and nucleolin (D4C7O) rabbit mAb (Cell Signalling cat # 14574) appropriately diluted in Odyssey^®^ Blocking Buffer (TBS) 0.1% TWEEN^®^ 20 (Sigma Aldrich cat # P1379) overnight at 4°C. The following day, the membrane was washed four times with 1xTBS 0.1% Tween-20 for 5min and then incubated with goat-anti mouse IgG IRDye680^®^ (Rockland cat # 610144002) and goat anti-rabbit IgG IRDye800^®^ (Rockland cat # 611132122) in Odyssey Blocking Buffer 0.1% Tween-20 for 1.5h at room temp. The membrane was washed as previously described and then dried completely before being scanned on Odyssey^®^ CLx Imaging System (LI-COR). Fluorescence was quantified using the Image Studio™ Lite (LI-COR) software.

### Subcellular fractionation and quantitative real-time PCR

Three independent wells from a 6-well culture plate (Sigma Aldrich cat # CLS3516) containing 1×10^6^ low passage HAP1 cells were lifted using 0.25% Trypsin-EDTA (Gibco cat # 25300096) and washed once with DPBS (Gibco cat # 14190144). Nuclear and cytoplasmic lysates were separated from cells and RNA isolated from both fractions using the PARIS™ Kit (Thermo Fisher cat # AM1921) following manufacturer’s instructions. RNA was treated with the TURBO DNA-free™ Kit (Life Technologies cat # AM1907) and concentration was measured on the NanoDrop™ Lite Spectrophotometer (Thermo Fisher Scientific cat # ND-LITE-PR). A qRT-PCR was performed on equal concentrations of DNA-free RNA from both fractions using the *Power* SYBR^®^ Green RNA-to-CT™ 1-Step Kit (Thermo Fisher Scientific cat # 4389986) with primers for MALAT1 (nuclear control), ACTB (cytoplasmic control) and PDCL3P4. Primers for PDCL3P4 were designed to avoid cross-detection of parent gene cDNA. The primers span an exon-exon junction whereby the forward primer is located in a novel upstream exon and the reverse primer in a novel extended sequence of the retroposed portion of PDCL3P4. Samples were run on the ViiA™ 7 Real-Time PCR machine following kit instructions and the ratio of nuclear and cytoplasmic expression was calculated where, nuclear enrichment= 2^Raw Ct Cytoplasm – Raw Ct Nucleus^ and cytoplasmic enrichment= 2^Raw Ct Nucleus – Raw Ct cytoplasm^. Ct values were not normalised to the differential abundance of the MALAT1 and ACTB housekeeping genes between cellular compartments.

## Ethics approval and consent to participate

Not applicable.

## Availability of data and materials

All PacBio data generated by this study is deposited in the Gene Expression Omnibus under accession GSE160383.

## Competing interests

The authors declare that they have no competing interests.

## Funding

This study was funded by the Australian Department of Health Medical Frontiers Future Fund (MRFF) (MRF1175457 to A.D.E.), the Australian National Health and Medical Research Council (NHMRC) (GNT1173711 to G.J.F. and GNT1161832 to S.W.C.), a CSL Centenary Fellowship to G.J.F., a University of Queensland Early Career Researcher Grant to S.W.C and by the Mater Foundation (Equity Trustees / AE Hingeley Trust).

## Authors’ contributions

G.J.F and S.W.C designed the study. R-.L.T, Y.J and S.W.C performed the experiments. R.-L. T, T.R.M, A.D.E, and S.W.C performed the analysis. S.W.C and G.J.F funded the study. R-.L.T, A.D.E, G.J.F and S.W.C wrote the manuscript. All authors read and approved the final manuscript.

## Acknowledgements

The authors would like to thank C. James for technical assistance, and acknowledge the Translational Research Institute (TRI) for research space and equipment that enabled this research. We would particularly like to thank the TRI flow cytometry core facility for assistance with this study. The authors thank the University of Queensland Genome Innovation Hub for continuing support.

## Supplementary figure legends

**Fig. S1.**
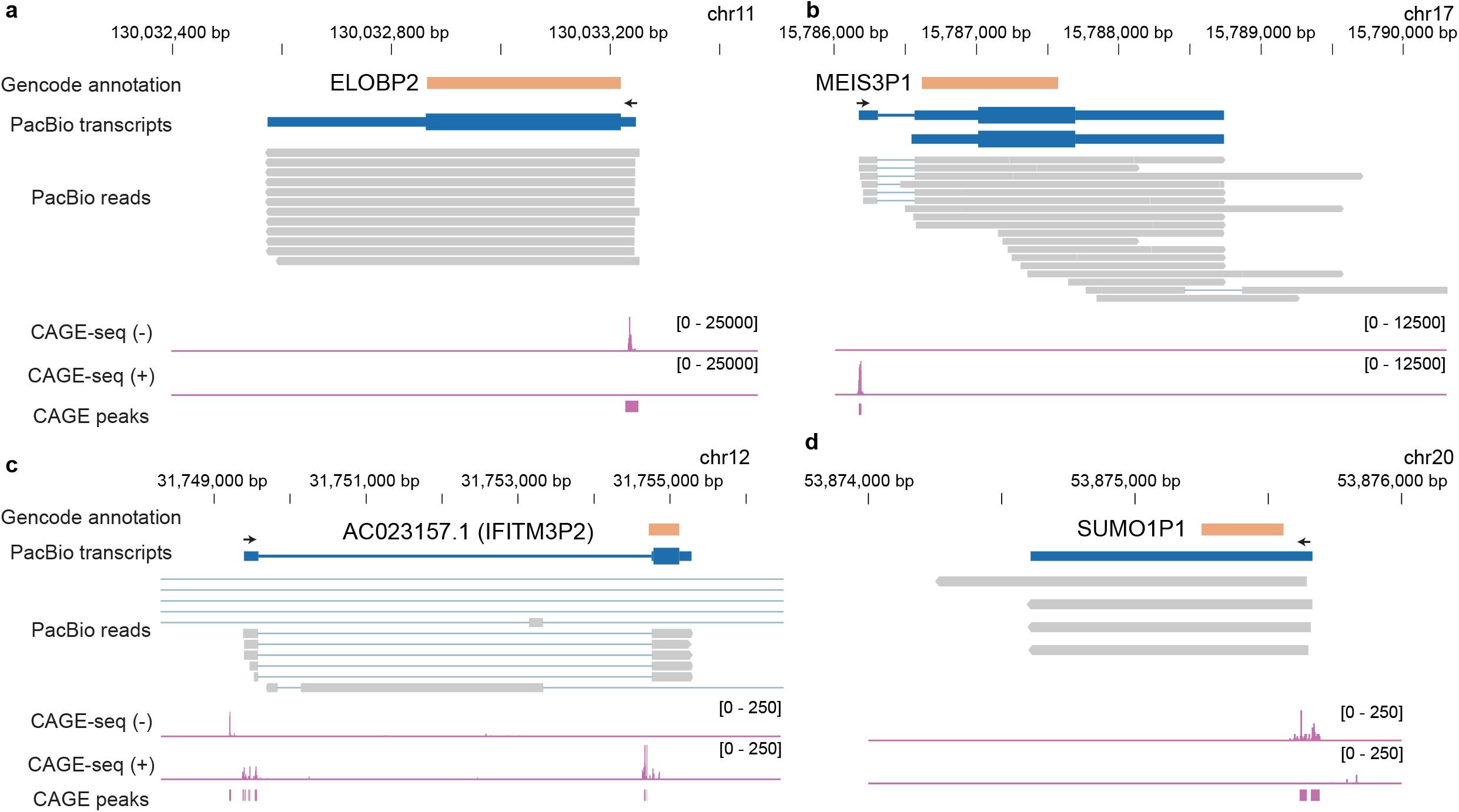
**a** PacBio Iso-Seq reads identify the full-length ELOBP2 transcript. The TSS of ELOBP2 is supported by a CAGE peak. **b** PacBio Iso-Seq reads identify the full-length MEIS3P1 transcript. The TSS of MEIS3P1 is supported by a CAGE peak. **c** PacBio Iso-Seq reads identify the full-length IFITM3P2 transcript. The TSS of IFITM3P2 is supported by a CAGE peak. **d** PacBio Iso-Seq reads identify the full-length SUMO1P1 transcript. The TSS of SUMO1P1 is supported by a CAGE peak.

**Fig. S2.**
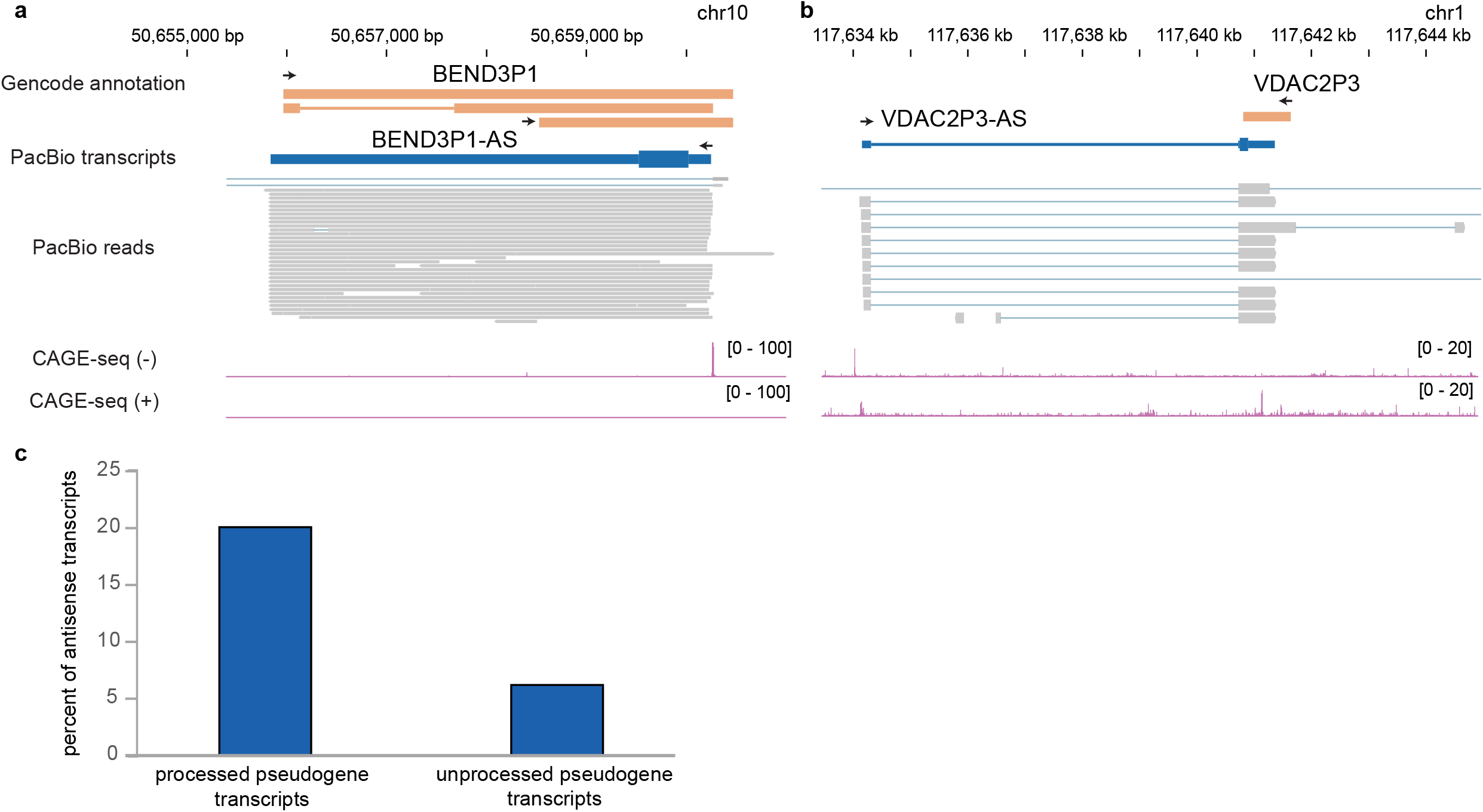
**a** BEND3P1 is expressed in the antisense orientation with respect to its parent gene. **b** The spliced pseudogene transcript VDAC2P3 is expressed in the antisense orientation with respect to its parent gene. **c** A higher fraction of processed pseudogenes transcripts are antisense compared to unprocessed pseudogenes.

**Fig. S3.**
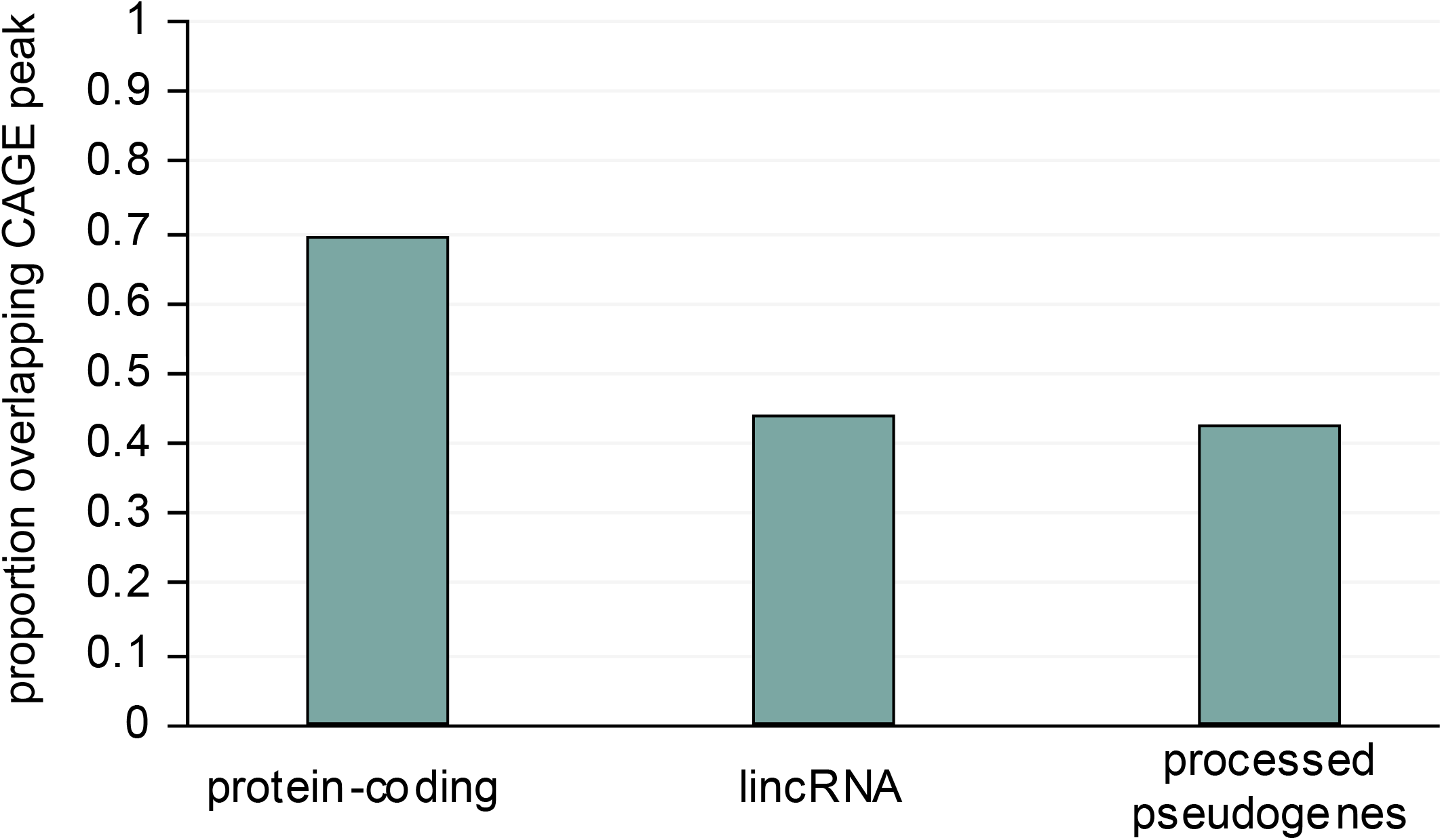
Proportion of transcripts of different biotypes overlapping with FANTOM5 CAGE peaks.

**Fig. S4.**
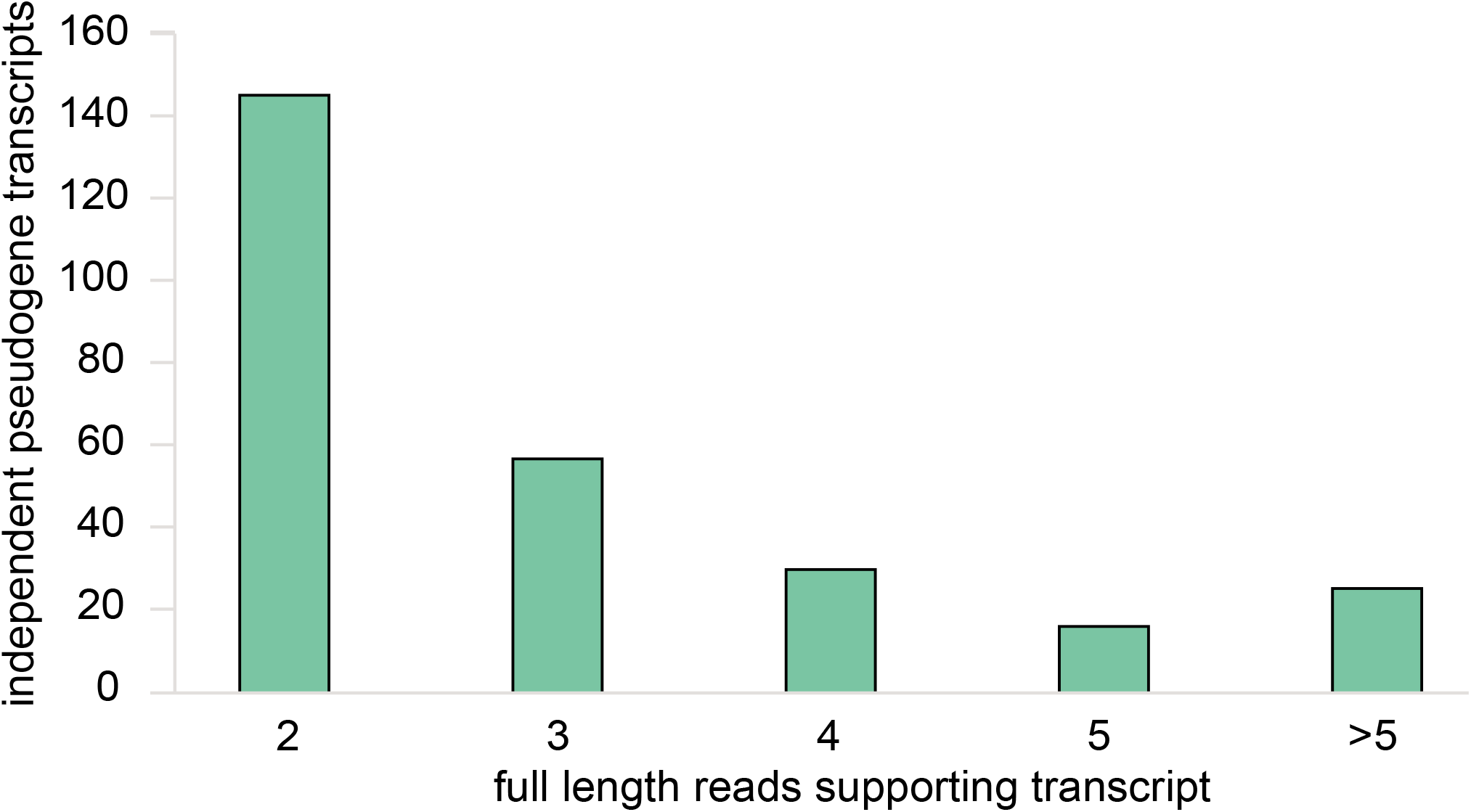
Distribution of number of full-length reads supporting each pseudogene transcript model. The minimum required reads is two.

**Fig. S5.**
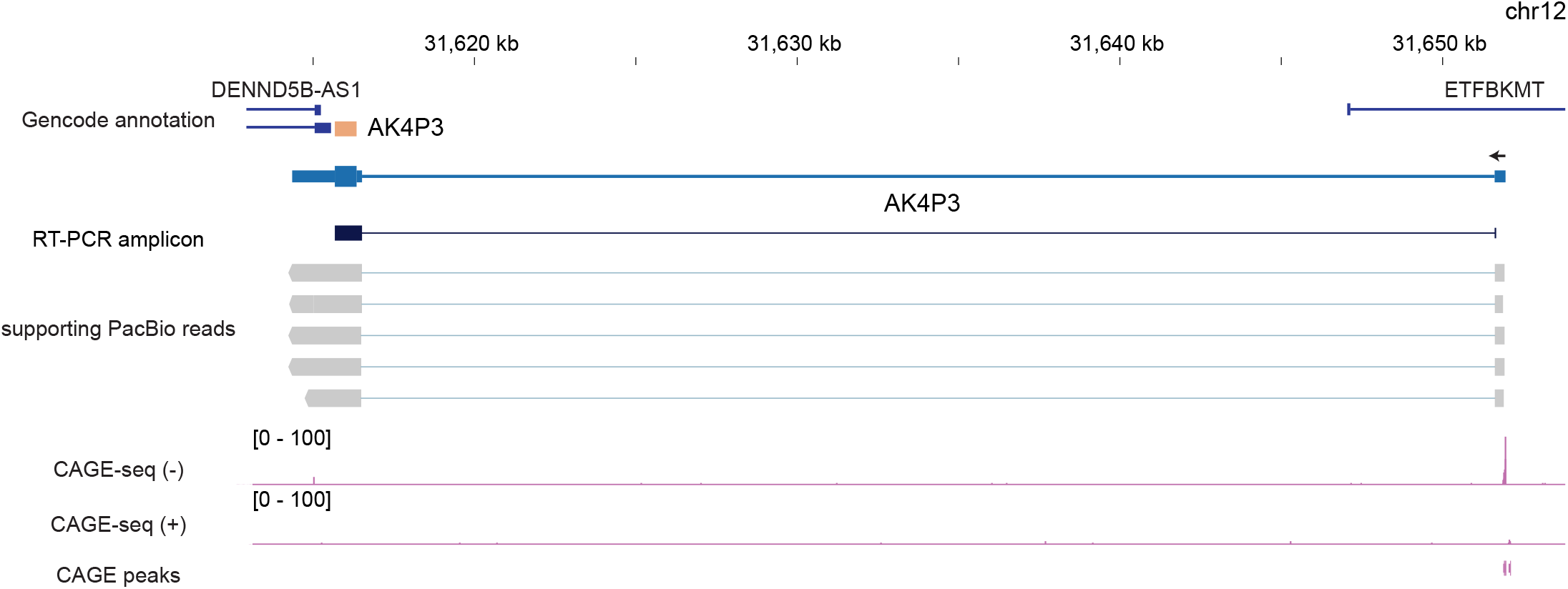
AK4P3 a transcribed potentially-coding pseudogene. AK4P3 has a novel 5’ exon and is transcribed from an upstream CAGE-confirmed TSS.

**Fig. S6.**
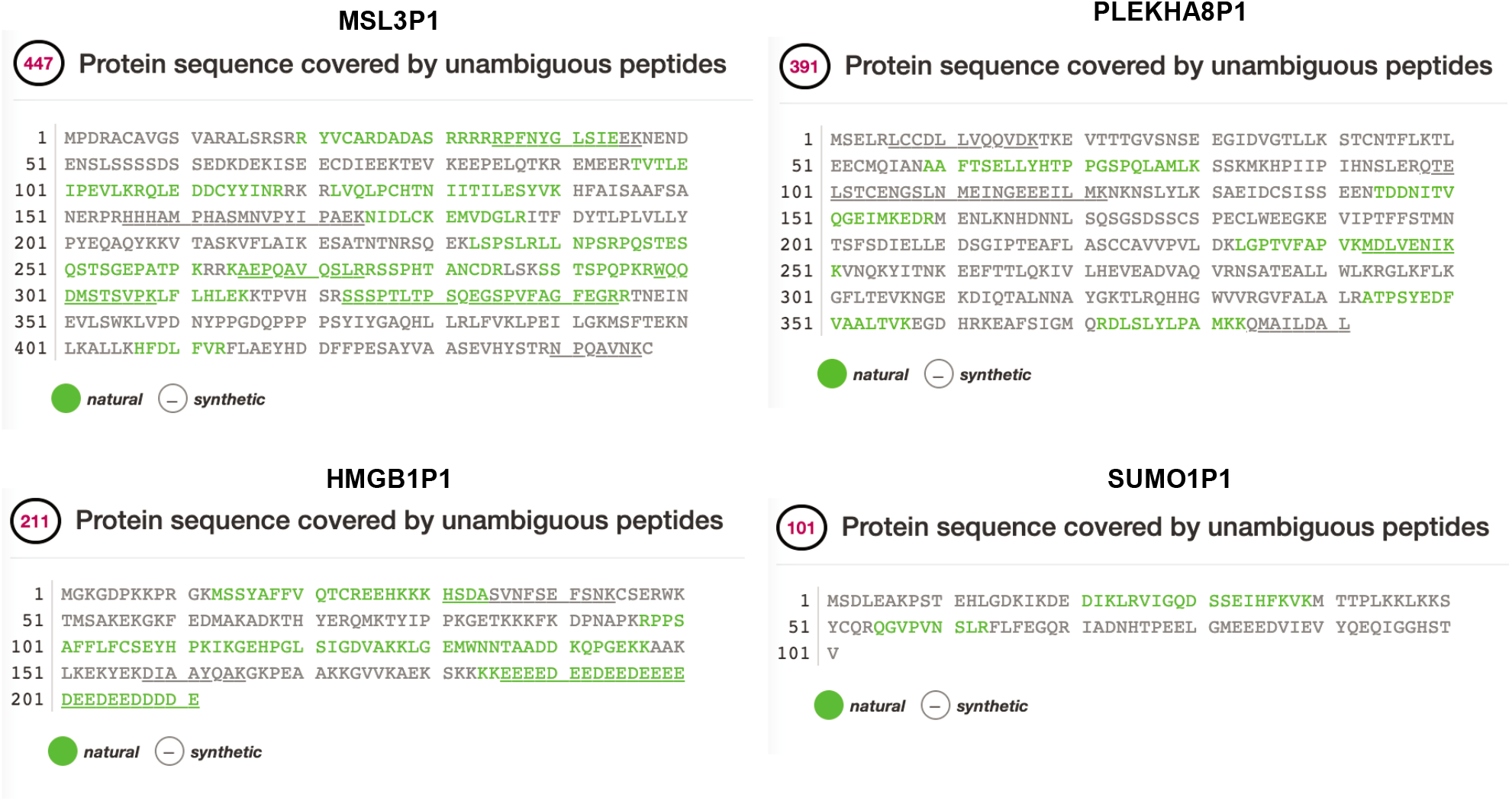
neXtprot peptide coverage of four translated pseudogenes. Natural peptides are those observed in mass-spectrometry experiments, whilst synthetic peptides are artificial standards.

**Fig. S7.**
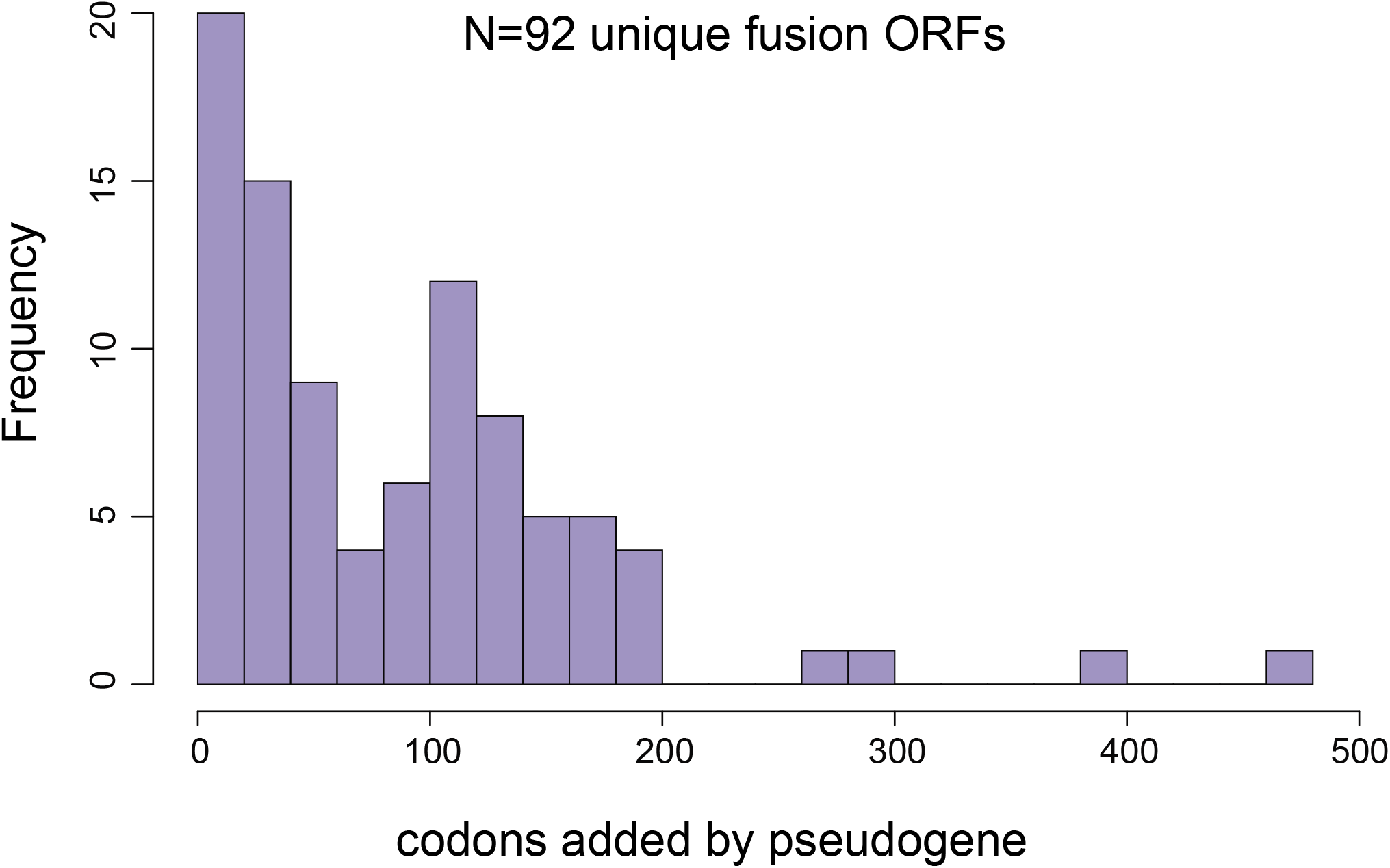
Distribution of the number of codons that pseudogenes contribute to known protein-coding genes.

**Fig. S8.**
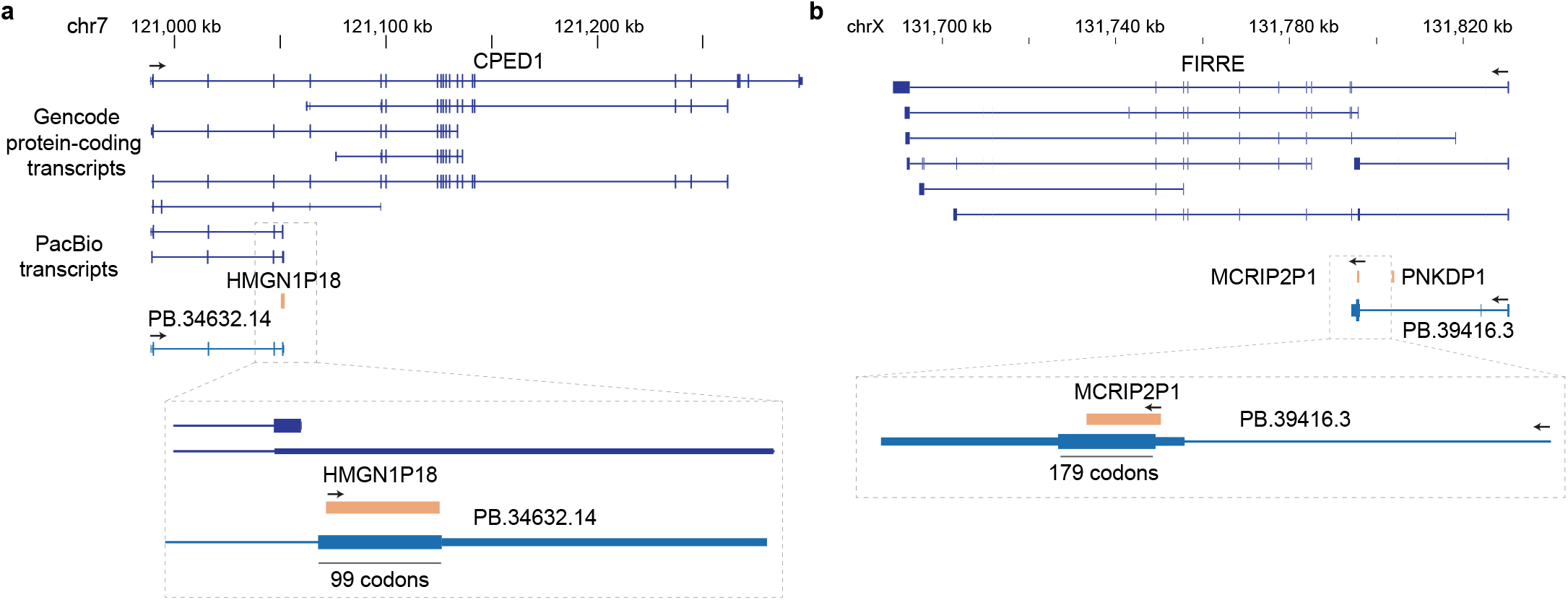
A novel CPED1 isoform contains a HMGN domain encoded by the pseudogene HMGN1P18. **b** An isoform of the lncRNA FIRRE splices into MCRIP2P1 and may encode a protein.

**Fig. S9.**
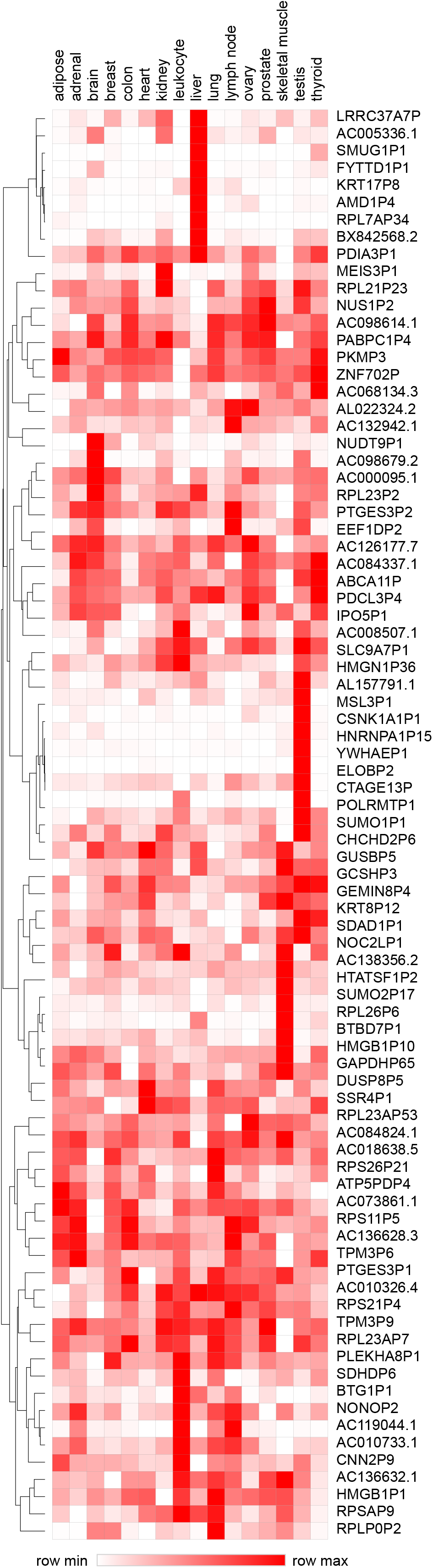
Tissue-specific pseudogene expression in the 16 adult tissues of the Illumina Body Map. Heatmap was generated using Morpheus, https://software.broadinstitute.org/morpheus. Expression is represented as log2(cpm+1) and is row scaled. Only sense independent pseudogenes with expression of greater than one cpm in at least one tissue are shown.

**Fig. S10.**
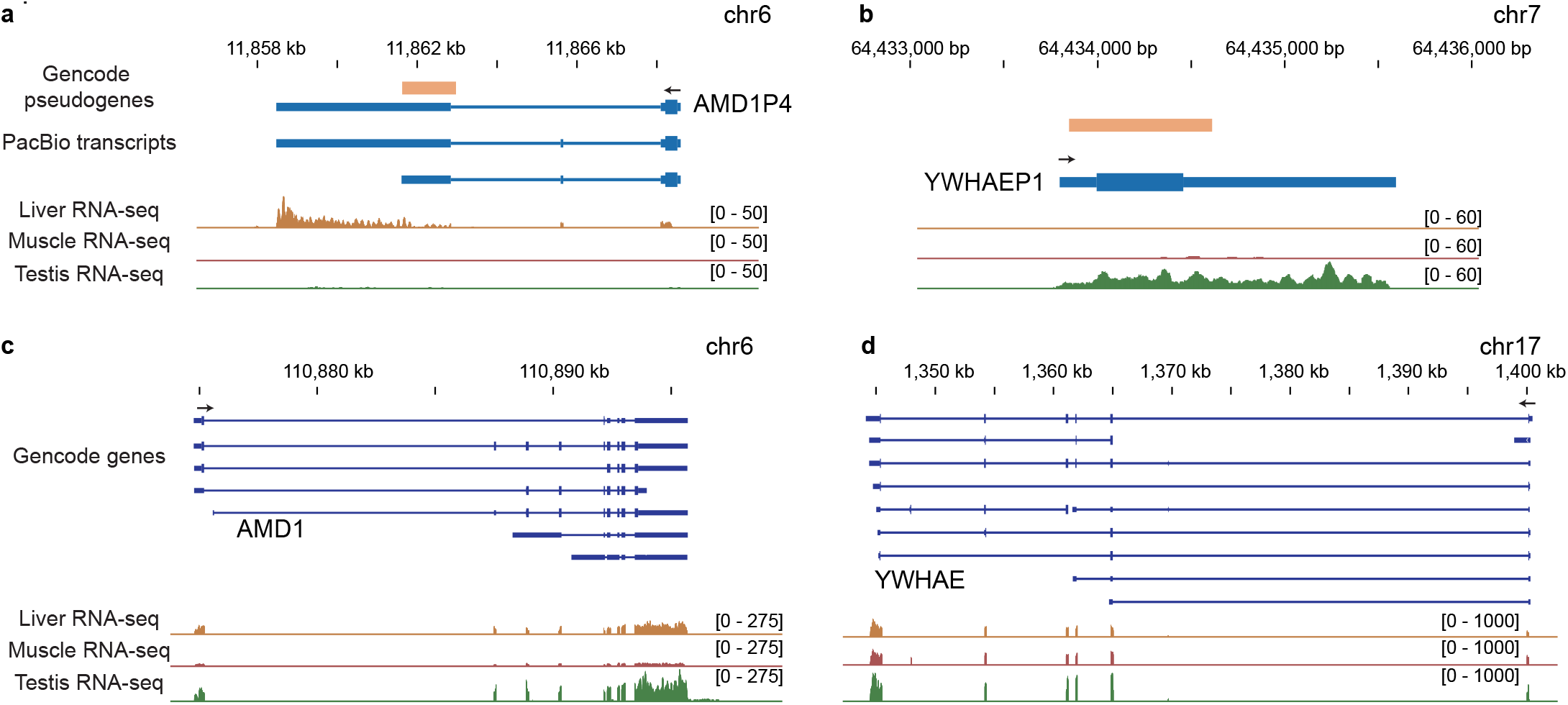
**a** AMD1P4 is expressed exclusively in the liver. **b** YWHAEP1 is expressed exclusively in the testis. **c** AMD1 is expressed broadly. **d** YWHAE is expressed broadly.

**Fig. S11.**
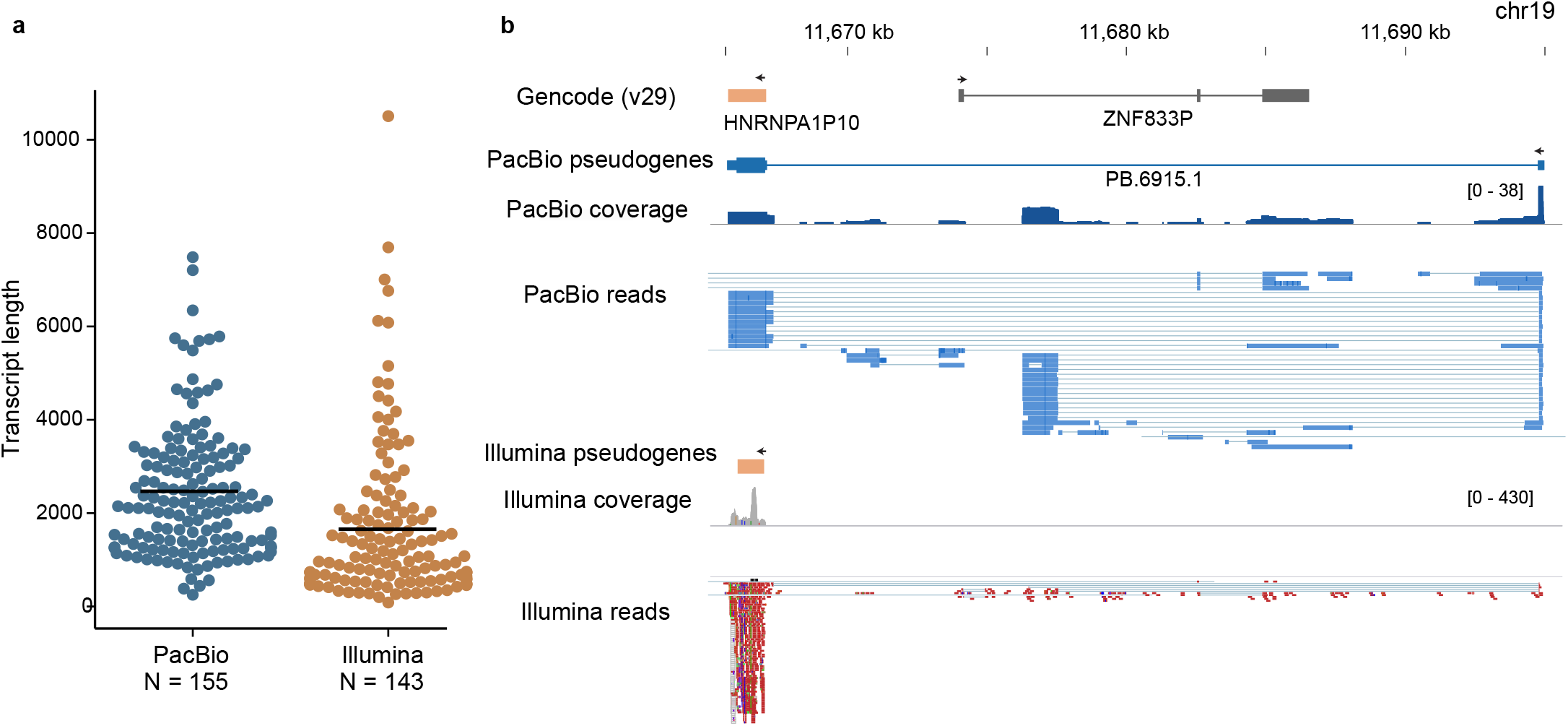
**a** Average of length of pseudogene transcripts assembled by StringTie [30] from Illumina data compared to PacBio pseudogene transcripts. Plot generated with Estimation Stats [58] **b** PacBio sequencing identifies a HNRNPA1P10 isoform that is not detected by short-read assembly.

**Fig. S12.**
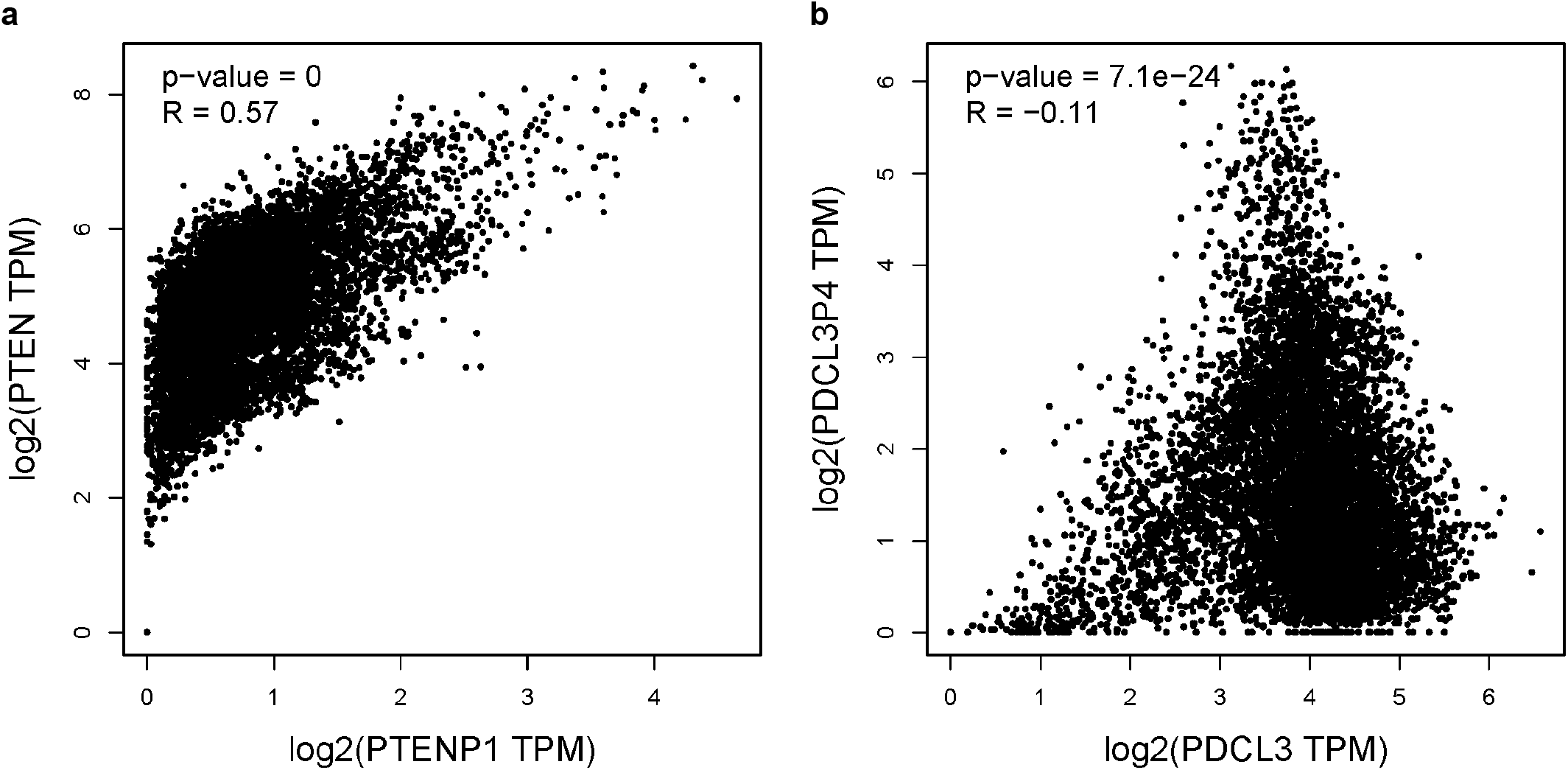
**a** PTEN and PTENP1 are strongly correlated in GTEX tissues. Correlation plots were generated with GEPIA [59]. **b** PDCL3P4 and PDCL3 expression is not positively correlated in GTEX [60] tissues.

## Supplementary tables

**Table S1** Characterisation of pseudogene transcripts identified by long-read sequencing.

**Table S2** Conservation of pseudogene ORFs across primate evolution.

**Table S3** Tissue-specific expression of pseudogene transcripts in the Illumina Body Map.

**Table S4** RNA-seq of PDCL3P4 knockout lines.

**Table S5** Oligonucleotides used in this study.

## Notes

### Competing Interest Statement

The authors have declared no competing interest.

